# Cooperative multivalent receptor binding promotes exposure of the SARS-CoV-2 fusion machinery core

**DOI:** 10.1101/2021.05.24.445443

**Authors:** Alexander J. Pak, Alvin Yu, Zunlong Ke, John A. G. Briggs, Gregory A. Voth

## Abstract

The molecular events that permit the spike glycoprotein of severe acute respiratory syndrome coronavirus 2 (SARS-CoV-2) to bind, fuse, and enter cells are important to understand for both fundamental and therapeutic reasons. Spike proteins consist of S1 and S2 domains, which recognize angiotensin-converting enzyme 2 (ACE2) receptors and contain the viral fusion machinery, respectively. Ostensibly, the binding of spike trimers to ACE2 receptors promotes the preparation of the fusion machinery by dissociation of the S1 domains. We report the development of bottom-up coarse-grained (CG) models validated with cryo-electron tomography (cryo-ET) data, and the use of CG molecular dynamics simulations to investigate the dynamical mechanisms involved in viral binding and exposure of the S2 trimeric core. We show that spike trimers cooperatively bind to multiple ACE2 dimers at virion-cell interfaces. The multivalent interaction cyclically and processively induces S1 dissociation, thereby exposing the S2 core containing the fusion machinery. Our simulations thus reveal an important concerted interaction between spike trimers and ACE2 dimers that primes the virus for membrane fusion and entry.

## Introduction

Infection by severe acute respiratory syndrome coronavirus 2 (SARS-CoV-2) begins with binding of viral particles to angiotensin-converting enzyme 2 (ACE2) cell-surface receptors, followed by viral-host cell membrane fusion, and entry of the viral genetic cargo (Shang et al., 2020). These events are orchestrated by spike glycoproteins – trimeric class I fusion proteins – that decorate the exterior of SARS-CoV-2 particles. Each spike protein consists of S1 and S2 domains, which are post-translationally cleaved by furin and remain non-covalently associated in a metastable prefusion state (Walls et al., 2020; Wrapp et al., 2020). The S1 domain contains the receptor binding domain (RBD) that is responsible for ACE2 recognition, while the S2 domain contains the fusion machinery. The S2 domains interact as a trimeric core with a characteristic helical stalk that is partially covered by the three S1 domains; the RBDs interact as a trimeric cap around the S2 trimeric core (Cai et al., 2020; Ke et al., 2020). A hinge-like conformational change exposes the RBD interfaces that are amenable to ACE2 association. Receptor binding is believed to result in S1 shedding, thereby exposing the S2 core. For fusion to occur, the second proteolytic site S2’ is cleaved, thereby releasing the fusion peptide, after which the S2 core undergoes a dramatic conformational change into an extended helical stalk, i.e., the postfusion state, in a process similar to prior coronaviruses (Walls et al., 2017).

As spike proteins are indispensable to SARS-CoV-2 infectivity, inhibiting their function is a natural target for therapeutic design. Understanding how spike proteins bind to ACE2 and how binding primes exposure of the S2 trimeric core is therefore essential to viral activity. Structures resolved using cryo-electron microscopy (cryo-EM) combined with docking have shown that ACE2-RBD binding requires spike trimers to be open, and that two spike trimers can be accommodated on one ACE2 dimer (Yan et al., 2020). Additional cryo-EM structures have shown that up to three soluble ACE2 monomers can bind to the same spike trimer, which may result in exposure of the S2 trimeric core (Benton et al., 2020); indeed, soluble ACE2 trimers were recently demonstrated as an effective inhibitor of SARS-CoV-2 (Xiao et al., 2021). Both modalities are potentially important for viral avidity, and furthermore, likely dependent upon the flexibility exhibited by both the spike trimer stalk (Turoňová et al., 2020) and ACE2 domain (Barros et al.), as well as the stochastic nature of RBD opening observed throughout the spike trimer (Cai et al., 2020; Ke et al., 2020). It has also been suggested that ACE2 binding to spike trimers induces allosteric effects that heighten (dampen) conformational motions near proteolytic cleavage sites and the fusion peptide (near the central helix and stalk) (Raghuvamsi et al., 2021) or conformational motions that promote subsequent RBD opening (Xu et al., 2021). Characterization of three circulating variant strains (B.1.1.7, B.1.351, and P.1) has shown that increased infectivity is, in part, due to mutations in the RBD that increase ACE2 binding affinity (Supasa et al., 2021; D. Zhou et al., 2021). Clearly, viral binding, fusion, and entry rely on a complex dynamical process that likely leverages the concerted interactions between spike trimers and ACE2 receptors.

To elucidate the molecular events that lead to binding and priming of the spike protein for membrane fusion, we use molecular dynamics simulations. We first use atomistic molecular dynamics (MD) simulations to systematically derive from “bottom-up” effective coarse-grained (CG) models of spike trimers and ACE2 dimers, which are then validated against conformational state statistics observed from experimental cryo-electron tomography (cryo-ET) data. We then use CG MD simulations at the larger scale of multi-protein arrays on membranes to investigate the binding between membrane-bound spike trimers and membrane-bound ACE2 receptors, and associated shedding of the S1 domains that expose the S2 trimeric cores. Our simulations show that multiple ACE2 dimers bind to spike trimers, consistent with prior cryo-EM models using soluble ACE2 (Benton et al., 2020). Our analysis indicates that binding between spike trimers and ACE2 dimers is positively cooperative, and furthermore, shows that this multivalent interaction is necessary to productively prime the virus for membrane fusion. At present, such insights are difficult to obtain experimentally.

## Results

Our CG MD simulations consist of three constituents, spike trimers, ACE2 dimers, and lipid membranes, each depicted in **Fig. 1A**. We extended our previously reported implicit-solvent, CG spike protein model derived from all-atom molecular dynamics trajectories (Yu et al., 2020), which were based on the open and closed state atomic models from cryo-EM (Protein Data Bank, PDB: 6VYB, 6VXX) (Walls et al., 2020). Each spike protomer is modeled as non-covalently interacting S1 and S2 domains with a CG resolution of around 10-15 amino acids per CG “bead” or site, and initially prepared as trimers. Briefly, intra-domain interactions are represented by a heterogeneous effective harmonic bond network while inter-domain interactions are represented by a combination of excluded volume repulsion, short-range attraction, and screened electrostatic terms (see *Materials and Methods*). A similar resolution and interaction model are used to represent each ACE2 monomer, initially prepared as dimers. The lipids are represented by a phenomenological CG model with a bending rigidity of 30 *k*_*B*_*T* to emulate the mechanical properties of typical membranes (Grime & Madsen, 2019). All CG model details and parametrization procedures are described in the *Materials and Methods* section.

**Fig. 1.**
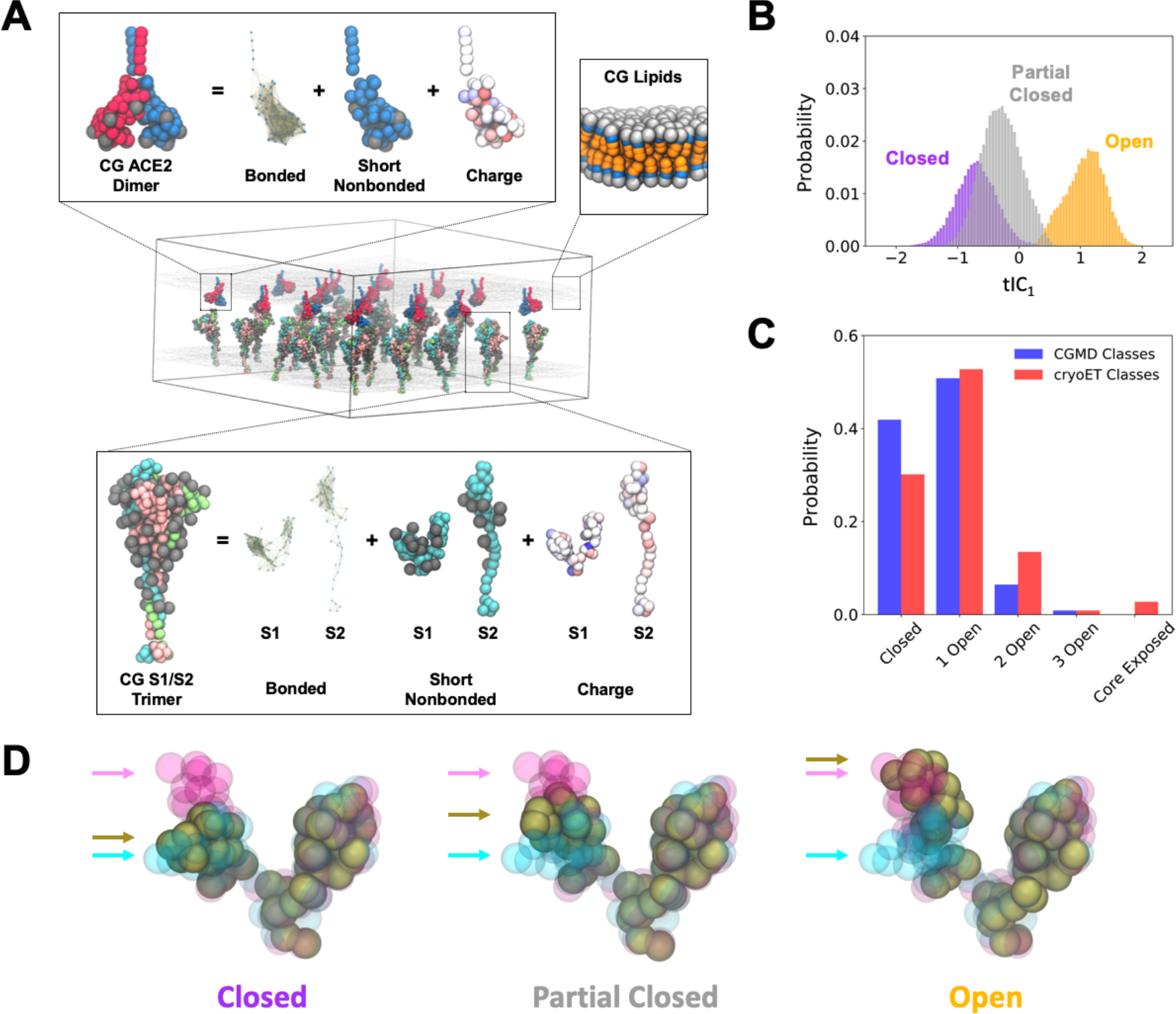
Schematic overview and characterization of our coarse-grained (CG) models. **(A)** Representative depiction of a CG simulation of spike trimers in membrane interacting with an adjacent membrane with ACE2 dimers. The insets depict the CG model components for the spike trimer (bottom), ACE2 dimer (upper left), and lipid membrane (upper right). Note that the protein CG sites are colored by monomer while glycans are represented by grey balls. **(B)** Probability distributions for RBD conformations projected onto the first time-lagged independent component (tIC_1_). The purple, grey, and orange distributions denote the three clusters identified by k-means clustering with labels indicating their representative conformational state. **(C)** Comparison of spike trimer conformational state populations (ranging from 0 to 3 open RBDs or shed RBDs, i.e., exposure of the S2 trimeric core) from CG simulations and cryo-ET classification; the core exposed state is in the prefusion form in the CG simulations while it is in the postfusion form in the cryo-ET dataset. **(D)** Representative depiction of the average configuration of CG S1 (brown beads) within the identified k-means clusters. For reference, CG configurations of experimentally-resolved open and closed states of S1 are shown as magenta and cyan beads, respectively. Arrows indicate the positions of each respective RBDs. In all cases, the N-terminal domains of S1 are aligned.

It is well-known that the RBDs of spike trimers are capable of hinge-like movements, resulting in combinations of open and closed configurational states throughout the trimer (Cai et al., 2020; Ke et al., 2020; Walls et al., 2020; Wrapp et al., 2020). Therefore, to validate our updated CG spike model, we quantified the configurational sampling of the RBD. We first simulated a single CG spike trimer in membrane (see *Materials and Methods*) for 50×10^6^ CG timesteps (*τ*_CG_ = 50 fs). To characterize the motions of the RBD, we quantified the distribution of reciprocal interatomic distances (T. Zhou & Caflisch, 2012) (DRID) for each configuration of the three RBDs; the DRID method computes the first, second, and third moments of the reciprocal distances between each CG site and every other CG site, yielding 3*N*_*CG*_ features. We next performed dimensional reduction using time-lagged independent component analysis (tICA), a linear projection technique that aims to maximize representation of auto-correlations (Naritomi & Fuchigami, 2011). The first 10 tIC dimensions were clustered using *k*-means clustering (Lloyd, 1982), revealing three primary conformational states. These three states can be seen in **Fig. 1B**, which depicts the probability distribution of each of the three labeled clusters along the first tIC (tIC_1_) dimension. Assessing the root-mean-squared deviation of the configurations from each of the three states (see **Fig. 1D and Fig. 1 – figure supplement 2**) reveals that the green-labeled distribution represents “closed” RBDs, the red-labeled distribution represents “open” RBDs, and the blue-labeled distribution represents an intermediate state we denote as “partially closed,” which we discuss later. The separability of the open- and closed-state distributions (we include the partially-closed state in the latter) depicted in **Fig. 1B** suggests that tIC_1_ can be used as a classifying metric for the open state (tIC_1_ > 0.4) and closed or partially-closed states (tIC_1_ < 0.4).

In our prior cryo-ET classification of RBDs within spike trimers along the surface of intact virions, we found that RBD conformations are stochastic and may range from three closed RBDs to three open RBDs, with three closed RBDs and one open (two closed) RBDs representing the two most predominant states (Ke et al., 2020). Given these observations, we assessed the distribution of open RBDs within spike trimers as predicted by our CG model. We prepared 25 spike trimers in a flat membrane (around 170×170 nm^2^) with periodic boundary conditions and simulated three replicas of the system for 172×10^6^ τ_CG_. The simulated surface density, which is around 1150 nm^2^ per spike trimer, was chosen to reflect virions observed using cryo-ET, which had 24 ± 9 spike trimers and an average diameter of 91 nm (Ke et al., 2020). Using the final 12.5×10^6^ τ_CG_, we characterized the RBDs throughout each of the trimers and computed the population of trimers with 0, 1, 2, and 3 open RBDs. In **Fig. 1C**, we compare the distribution of trimer states from our CG simulations to that of a previously reported cryo-ET dataset (Ke et al., 2020). Our CG spike trimers undergo a slow yet reversible transition between open and closed states, with around 50% of the total population having 1 open RBD, around 40% having 0 open RBDs (i.e., 3 closed), and diminishing populations of 2 and 3 open RBD trimers. The relative populations of these states are consistent with the cryo-ET classification of RBDs in virions. Note that complete S2 trimeric core exposure (i.e., the precursor to the postfusion state) was not observed throughout these simulations. However, we observed partial exposure in rare cases, i.e., single S1 domains from spike trimers spontaneously dissociated due to thermal fluctuations, which we discuss further later. In contrast, a minority population of postfusion trimers was observed by cryo-ET. At this point, it is unclear whether this discrepancy is due to limitations of the CG model, due to fixation by formaldehyde which may shift the open/closed equilibrium, or both. Nonetheless, our analysis provides qualitative validation that the conformational variability of the RBD in our CG spike trimer is consistent with experimental observations.

Next, to investigate the spike-ACE2 binding process, we simulated spike trimers interfacing with ACE2 dimers. We prepared triplicate simulations of spike trimers in one membrane adjacent to another containing ACE2 dimers (see **Fig. 1A**). The density of spike trimers and ACE2 dimers was fixed such that the relative ratio of ACE2 dimer to spike trimer ([ACE2_dimer_]/[S_trimer_]) was 2.56, i.e., 64 ACE2 dimers and 25 spike trimers, sufficient to allow multivalent ACE2 binding to each spike trimer. The two membranes (and their constituents) were initially shifted along their normal directions until the closest distance between spike trimers and ACE2 dimers was around 2 nm. The simulations were run for 172×10^6^ τ_CG_.

Throughout our simulations, we observed spike trimers fluctuating between the open and closed state, with the former preceding ACE2 binding, as expected. Interestingly, we observed spike trimers that were capable of binding to multiple ACE2 dimers, which required additional opening of RBDs. However, we did not see multiple spike trimers binding to the same ACE2 dimer, suggesting that any multivalent interactions between spike and ACE2 are due to an excess of accessible ACE2 receptors.

Our simulations predict two modalities for ACE2-facilitated dissociation of S1. First, S1 dissociation appears to occur due to thermal fluctuations, which may happen both in the absence and presence of binding between an open RBD and a single ACE2 dimer. The remaining two S1 monomers persist on the spike trimer, although in some cases, complete S1 shedding occurs after secondary RBD opening and binding to a new ACE2 receptor. The second modality was observed more frequently and involves the sequential binding of two ACE2 receptors to the same spike trimer. Consequently, dissociation of all three S1 domains is accelerated and results in the exposure of the S2 trimeric core (**Video 1**).

We depict the representative elements of the multivalent ACE2-binding process in **Fig. 2**, in addition to tIC1 time-series profiles for each RBD (additional examples are shown in **Fig. 2 – figure supplement 1**). Here, the observed effect on the bound spike trimer is one of processive release of S1 domains in a cyclical fashion. The process generally adheres to the following sequential steps. First, RBDs conformationally vary between the open and closed state, as seen for τ < 5×10^6^ τCG (note the variation of the cyan and pink lines/protomers in **Fig. 2**). One ACE2 dimer successfully binds to the open RBD, pulling the RBD and enforcing the open state, as seen for τ = 12×10^6^ CG (the cyan protomer adopts tIC_1_ = 1.0 with reduced variance). The cleft formed between the upraised RBD and the N-terminal arm of S1 provides a stabilizing pocket for the clockwise RBD (from a top-down view), or the green protomer as depicted in **Fig. 2** (tIC_1_ = −0.6), thereby “tightening” the adhesion of this S1 domain. Conversely, the counter-clockwise RBD, or the pink protomer as depicted in **Fig. 2**, appears to “loosen” its adhesion to the same cleft provided by the green S1 domain. Due to the weakened interactions, the pink protomer dynamically adopts the partially-closed state, as seen between 15×10^6^ < τ < 20×10^6^ τ_CG_ (pink line/protomer in **Fig. 2** with tIC_1_ = −0.25). Thermal fluctuations allow the pink protomer to spontaneously adopt the open state (τ = 26×10^6^ τ_CG_). A second ACE2 dimer binds to the newly exposed RBD and “pulls” this S1 domain away from the spike, thereby promoting dissociation (τ = 30×10^6^ τ_CG_). The remaining S1 dimer complex, which remains bound to the first ACE2 dimer, subsequently “loosens” and eventually dissociates due to thermal fluctuations (τ = 45×10^6^ τ_CG_). Finally, the S1-bound ACE2 dimers diffuse away from the spike trimer, which now consists of the exposed S2 trimeric core (τ > 50×10^6^ τ_CG_).

**Fig. 2.**
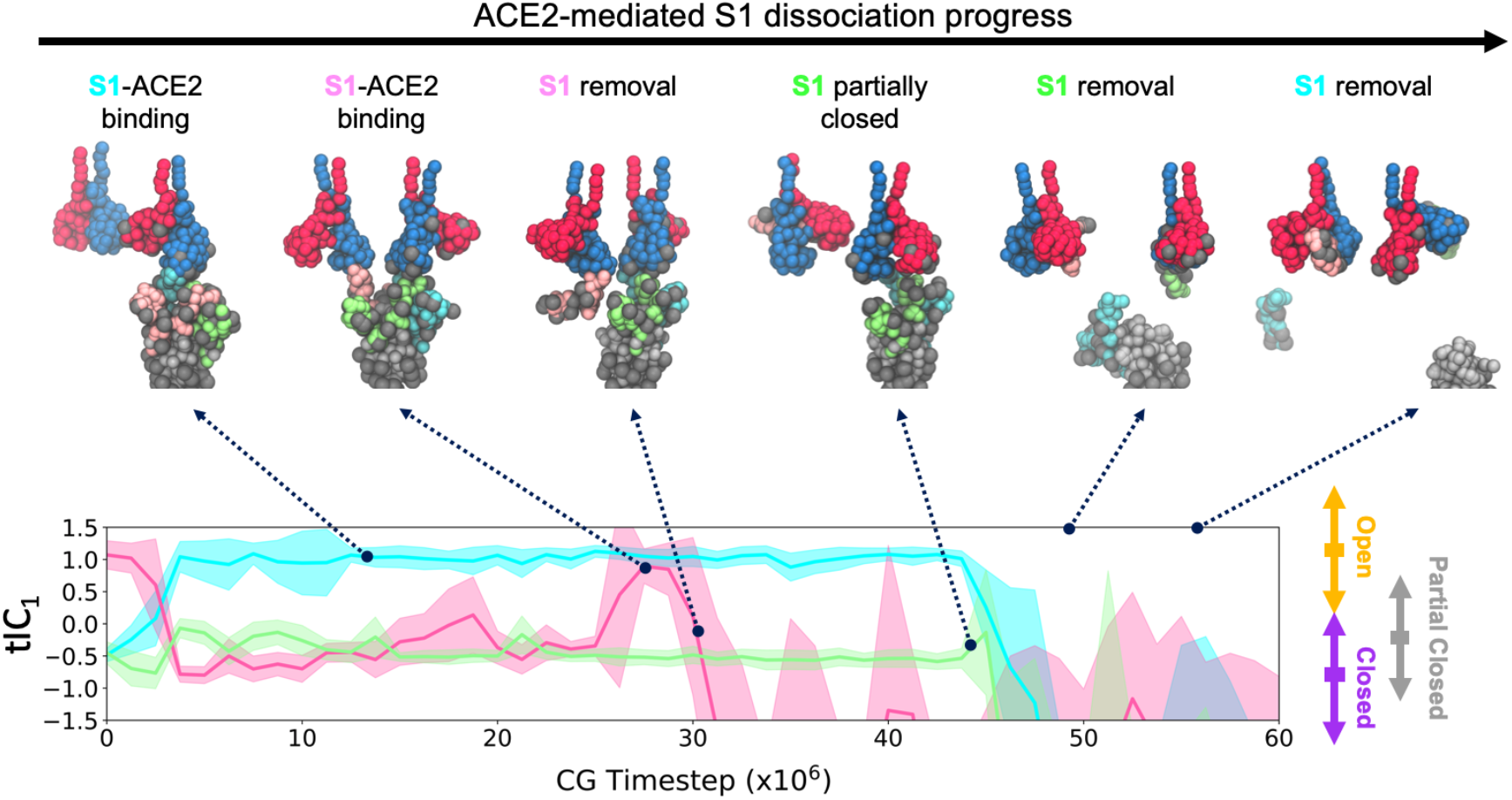
Dissociation of S1 facilitated by multivalent ACE2 binding. (Top) Snapshots of one representative S1 dissociation process upon binding of two ACE2 dimers. Each monomer in the ACE2 dimer is represented by red and blue beads, respectively, while each S1 protomer in the spike trimer is represented by cyan, pink, and green beads, respectively. The grey and silver beads denote glycans and S2 protomers, respectively. The depicted process is also shown in **Video 1. (Bottom)** Time series profile depicting tIC_1_ evaluated for the RBD of each protomer (sharing the same color) for the spike trimer depicted above, i.e., spike trimer protomers are labeled cyan to pink to green (back to cyan) in counter-clockwise order when viewing from the top-down. To the right of the panel, the double-ended arrows represent the extent of the tIC_1_ distributions for each state from **Fig. 1B**, while the rectangle shows the peak of each distribution.

Prior experimental binding assays using soluble ACE2 have demonstrated positive cooperativity when binding to SARS-CoV-2 spike trimers (Anand et al., 2020). Our simulations show that multivalent ACE2 interactions induce productive S1 dissociation, and further suggest that exposure of the postfusion machinery may be cooperatively correlated with ACE2 expression. To test this hypothesis, we prepared additional triplicate simulations with the same spike trimer density but varied the ACE2 density. Here, we tested stoichiometric ratios of [ACE2_dimer_]/[S_trimer_] between 0 and 4, the latter representing 100 ACE2 dimers and 25 spike trimers, and performed simulations for 172×10^6^ τ_CG_. The final 12.5×10^6^ τ_CG_ were used for analysis.

We summarize statistics on the extent of ACE2 binding, S1 dissociation, and S2 trimeric core exposure as [ACE2_dimer_]/[S_trimer_] is varied in **Fig. 3A**. We used a distance-based metric (described in *Materials and Methods*) to determine both binding and dissociation events. First, we find that S1 monomers are capable of spontaneous dissociation due to thermal fluctuations in the absence of ACE2, as noted above. Within our simulated timescales, however, none of these dissociation events resulted in complete S2 trimeric core exposure. As S1 dissociation was not observed in our atomistic MD simulations, nor during training of the CG model, we speculate that the collective membrane fluctuations, further induced by the spike trimer inclusions, promote S1 dissociation as bound S1 should be considered a metastable state.

**Fig. 3.**
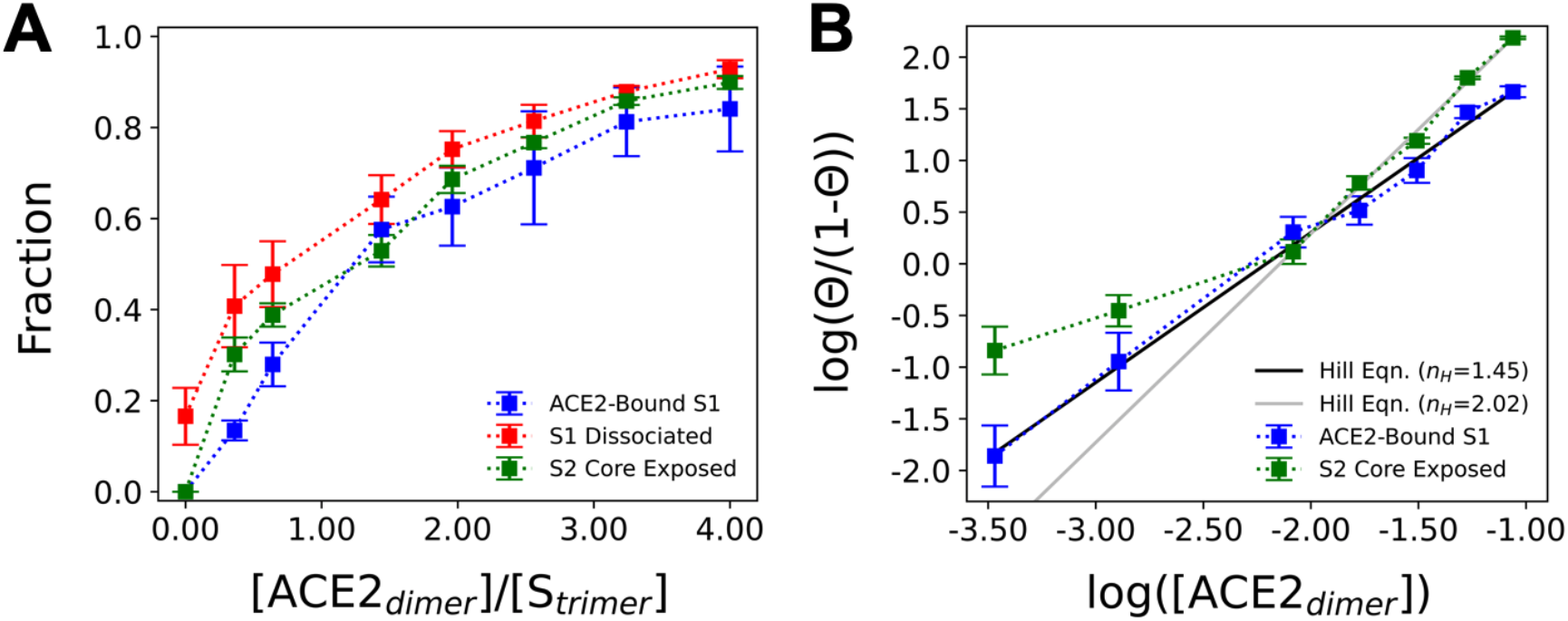
Cooperative enhancement of S2 trimeric core exposure by ACE2. **(A)** Summary statistics for the fraction of S1 monomers bound to ACE2 (blue), S1 monomers dissociated from the spike trimer (red), and spike trimers with complete exposure of the S2 trimeric core (green) as the stoichiometric ratio of ACE2 dimers to spike trimers, [ACE2_dimer_]/[S_trimer_], is varied. **(B)** Hill plot comparing the natural log of the surface density of ACE2 (in #/nm^2^) to the natural log of the fraction of S1 monomers bound to ACE2 or the fraction of completely exposed S2 trimeric cores (Θ). The black and grey lines are fits to the Hill equation with Hill coefficients (*n*_*H*_) equal to 1.45 and 2.02. All data points report the mean and standard error from triplicate simulations.

The presence of ACE2 has intriguing effects on binding, dissociation, and exposure statistics. To measure the degree of cooperativity, we fit our binding data to the Hill equation:

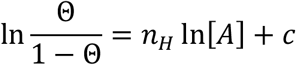

where Θ is the fraction of S1 monomers bound to ACE2, [*A*] is the surface density of ACE2, *n*_*H*_ is the Hill coefficient, and *c* is a constant. As seen in **Fig. 3B**, we find that *n*_*H*_ > 1 (= 1.45) for the binding curve, indicating that S1 binding to ACE2 is positively cooperative. In **Fig. 3A**, both the extent of S1 dissociation and S2 trimeric core exposure appear to monotonically increase with [ACE2_dimer_]/[S_trimer_]. Note that the relative ratio between S1 dissociation and S2 core exposure significantly decreases when [ACE2_dimer_]/[S_trimer_] ratios exceed 2. In fact, as seen in **Fig. 3B**, the propensity for S2 core exposure when [ACE2_dimer_]/[S_trimer_] > 2 also exhibits positive cooperativity with *n*_*H*_ = 2.02. Together, these results suggest that multivalent ACE2 binding promotes S1 dissociation events that productively expose the S2 trimeric core, and eventually, the postfusion machinery.

## Discussion

We developed and applied CG simulation models of SARS-CoV-2 spike trimers and receptor ACE2 dimers to study the dynamical mechanisms involved in viral avidity and dissociation, the latter being a necessary step prior to membrane fusion. Our simulations recapitulate the metastability of the S1 domains, which remain bound to spike trimers after furin cleavage. Although rare, we observed spontaneous dissociation of S1 even in the absence of ACE2 binding, consistent with cryo-EM characterization of postfusion trimers on mature virions (Cai et al., 2020; Ke et al., 2020). It has been shown that the D614G mutation is more infectious and stabilizes interactions between S1 and S2 that prevent dissociation (L. Zhang et al., 2020), promotes RBD opening (Benton et al., 2021; Mansbach et al., 2021), and increases ACE2 affinity (Ozono et al., 2021). Ostensibly, the D614G mutation prevents premature S1 dissociation and facilitates productive RBD-ACE2 binding, while ACE2-mediated S1 dissociation remains unimpeded. It would be valuable for future atomistic and CG simulations to ascertain how D614G and related mutations (e.g. the B.1.1.7, B.1.351, B.1.617.2, and P.1 variants) affect spike trimer stability before, during, and after ACE2 binding.

One of our key findings is that the primary binding modality between spike trimers and ACE2 dimers involves multiple ACE2 dimers binding to single spike trimers, rather than multiple spike trimers binding to the same ACE2 dimer, an observation that is consistent with split reporter assays of soluble spike trimers incubated with ACE2 dimers (Lui et al., 2020). We found that spike trimer binding exhibits positive cooperativity with ACE2 dimer density due to the multivalent nature of ACE2 binding. Soluble ACE2 has also been observed to cooperatively bind to SARS-CoV-2 spike trimers, yet interestingly, not for SARS-CoV-1 spike trimers (Anand et al., 2020). Finally, our simulations suggest that the multivalency of the ACE2 to spike trimer interaction is necessary to facilitate complete dissociation of the S1 monomers that occlude the S2 trimeric core, which contains the viral fusion machinery. Binding to ACE2 induces a cyclical tightening and loosening process that processively weakens S1 interactions, both to adjacent S1 monomers and to the underlying S2 core, which leads to S1 dissociation. On the basis of cryo-EM maps of soluble spike trimers bound to soluble ACE2, it was recently proposed that the three RBDs throughout the spike trimer open and bind to ACE2 in a processive fashion, eventually resulting in simultaneous dissociation of the three S1 monomers due to reduced contacts between neighboring monomers (Benton et al., 2020). The process we observe in our simulations is conceptually similar, although we propose that only two ACE2 dimers are minimally necessary for productive uncoating of S1, and the counter-clockwise S1 (from a top-down view) from the initial ACE2-bound S1 tends to open, bind to a second ACE2, and dissociate first.

One notable difference between the conditions of our simulations and the experiments discussed above is that both our spike trimers and ACE2 dimers are tethered to their respective membranes in a more realistic system than solubilized proteins. The tethers inherently restrict the diffusion and conformational motions available to both spike and ACE2 proteins. One potential consequence is that the relative populations of different spike/ACE2 complexes for solubilized proteins may not reflect populations at the interface between virions and host cells. Likewise, these populations are likely dependent upon the relative stoichiometry of spike trimers and ACE2 dimers, as our simulations indicate. We suggest that the use of fluid membrane-bound spike trimers and ACE2 receptors in biochemical and structural characterization of viral avidity, e.g., as seen in the work of Yang et al. and Lu et al. (Lu et al., 2020; Yang et al., 2020), compared to assays using soluble components, may prove to be characteristically different. For example, single-molecule Förster resonance energy transfer analysis suggests that spike trimers bound to one or two ACE2 receptors exhibit asymmetry throughout the three spike protomers (Lu et al., 2020), which is consistent with the dynamical behavior observed in our CG simulations.

Our analysis also motivates several directions for future simulations. While researchers have explored the free energy surface of single RBD open-close transitions (Casalino et al., 2021; Gur et al., 2020), the thermodynamics of secondary and tertiary RBD opening, as well as the role of multivalent ACE2 binding, should be thoroughly investigated. We note that our current CG model will also likely require additional features to holistically probe SARS-CoV-2 virion binding, fusion, and entry. Here, we investigated spike trimer and ACE2 dimer association at the interface of two planar membranes. However, it will be important to elucidate the effects of virion curvature (between 70-120 nm diameters (Ke et al., 2020)), which likely impose additional restrictions to spike trimer conformational sampling while seeking ACE2 receptors. Sophisticated modeling approaches to incorporate large-scale conformational changes, such as the extension of the heptad repeat coiled coil in the postfusion state (Walls et al., 2017), will also be necessary. In this work, we linearly mixed two model Hamiltonians that represent different configurational states (open and closed RBDs). Several additional approaches, including plastic network models (Maragakis & Karplus, 2005) and multi-configurational coarse-graining (Sharp, Vázquez, Wagner, Dannenhoffer-Lafage, & Voth, 2019), may prove useful to construct CG models capable of state transitions with increased accuracy for state populations and transition rates. These efforts will expand the capabilities of CG modeling, which in turn will provide valuable insight into collective molecular phenomena during SARS-CoV-2 infection, such as the role of cooperative multivalent ACE2 binding to induce spike trimer uncoating reported herein.

## Materials and Methods

### Atomistic MD simulations

Trajectories from prior atomistic MD simulations of the spike trimer in membrane were used to derive effective CG Hamiltonians (Casalino et al., 2020; Yu et al., 2020). Statistics for the open and closed state of the spike trimer were supplemented with additional triplicate simulations of the spike ectodomain using PDB 6VYB and 6VXX as initial structures, respectively (Walls et al., 2020). Triplicate simulations of the ACE2 dimer interacting with bound RBDs were also prepared using an initial structure from PDB 6M17 (Yan et al., 2020). All simulations were prepared using CHARMM-GUI (Jo, Kim, Iyer, & Im, 2008) and the CHARMM36m (Huang et al., 2016) force field. Each system was solvated with 2 nm of TIP3P water from the protein edge to the simulation domain boundary and with 150 mM NaCl. All simulations were performed using GROMACS 2020 (Abraham et al., 2015). Minimization was performed using steepest descent until the maximum force was reduced to 1,000 kJ/mol/nm^2^. Then, equilibration was performed in several phases. First, 10 ns were integrated in the constant *NVT* ensemble using the stochastic velocity rescaling thermostat (Bussi, Zykova-Timan, & Parrinello, 2009) with a damping time of 0.2 ps and a timestep of 1 fs. During this phase, the C*α* backbone of the protein was harmonically restrained with a force constant of 1,000 kJ/mol/nm^2^. An additional 40 ns were integrated in the constant *NPT* ensemble using the stochastic velocity rescaling thermostat (Bussi et al., 2009) (2 ps damping time) and the Parrinello-Rahman barostat (Parrinello & Rahman, 1980) (10 ps damping time) and a timestep of 2 fs. Finally, an additional 1650 ns were integrated in the constant *NVT* ensemble using the Nose-Hoover chain thermostat (Martyna, Klein, & Tuckerman, 1992) (2 ps damping time) and a timestep of 2 fs. Throughout this procedure, H-containing bonds were constrained using the LINCS algorithm (Hess, Bekker, Berendsen, & Fraaije, 1997). Statistics were gathered every 100 ps and the final 1,200 ns of each trajectory was used for CG model generation.

### CG modeling

CG model generation for the protein constituents proceeded in three phases. The first phase determined the mapping between atomistic and CG sites. Mapping was performed using the essential dynamics coarse-graining (EDCG) method (Z. Zhang et al., 2008), which was designed to preserve the principal modes of motion sampled during atomistic simulations (Z. Zhang et al., 2008). In EDCG, the CG mapping operator, 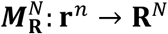, which relates the configurations of the atomistic trajectory (**r**^*n*^) to that of the CG model (**R**^*N*^), is variationally optimized using simulated annealing. Here, each CG site is constructed using the center of mass of residues within contiguous segments of the protein’s amino acid sequence. For *N* CG beads, *N* − 1 segment boundaries throughout the amino acid sequence are adjusted to minimize the target residual:

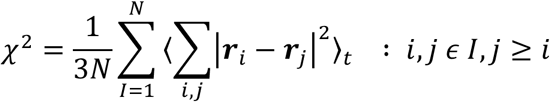

where *I* = 1, … , *N* is the CG site index, the brackets ⟨·⟩_*t*_ denote a time-averaged quantity, the sum over *i, j* is a sum over all unique pairs in the set of atoms belonging to the CG site *I*, and ***r***_*i*_ = **x**_*i*_ − ⟨ **x**_*i*_ ⟩_*t*_ is the displacement of atom *i* from its mean position ⟨ **x**_*i*_ ⟩_*t*_. Note that the residual is small when the displacements ***r***_*i*_ and ***r***_*j*_ are similar, i.e. the motions of atoms in the same CG site are correlated. We chose *N* using an “elbow” heuristic to identify when increasing *N* yielded diminishing returns in *χ*^2^. The “elbow” region when comparing a range of *N* values with corresponding residual (i.e., *χ*^2^) values, as depicted in **Fig. 4**, yields a CG mapping resolution that balances computational expense and least-squared error in the represented principal component subspace. Following this protocol, we used 60, 50, and 70 CG beads for the S1, S2, and ACE2 protein mappings, respectively. The N-linked glycans (Watanabe, Allen, Wrapp, McLellan, & Crispin, 2020) were removed prior to this procedure. Each glycan was mapped to its center of mass and appended to their corresponding EDCG-mapped protein, totaling 13, 9, and 7 sites, respectively.

**Fig. 4.**
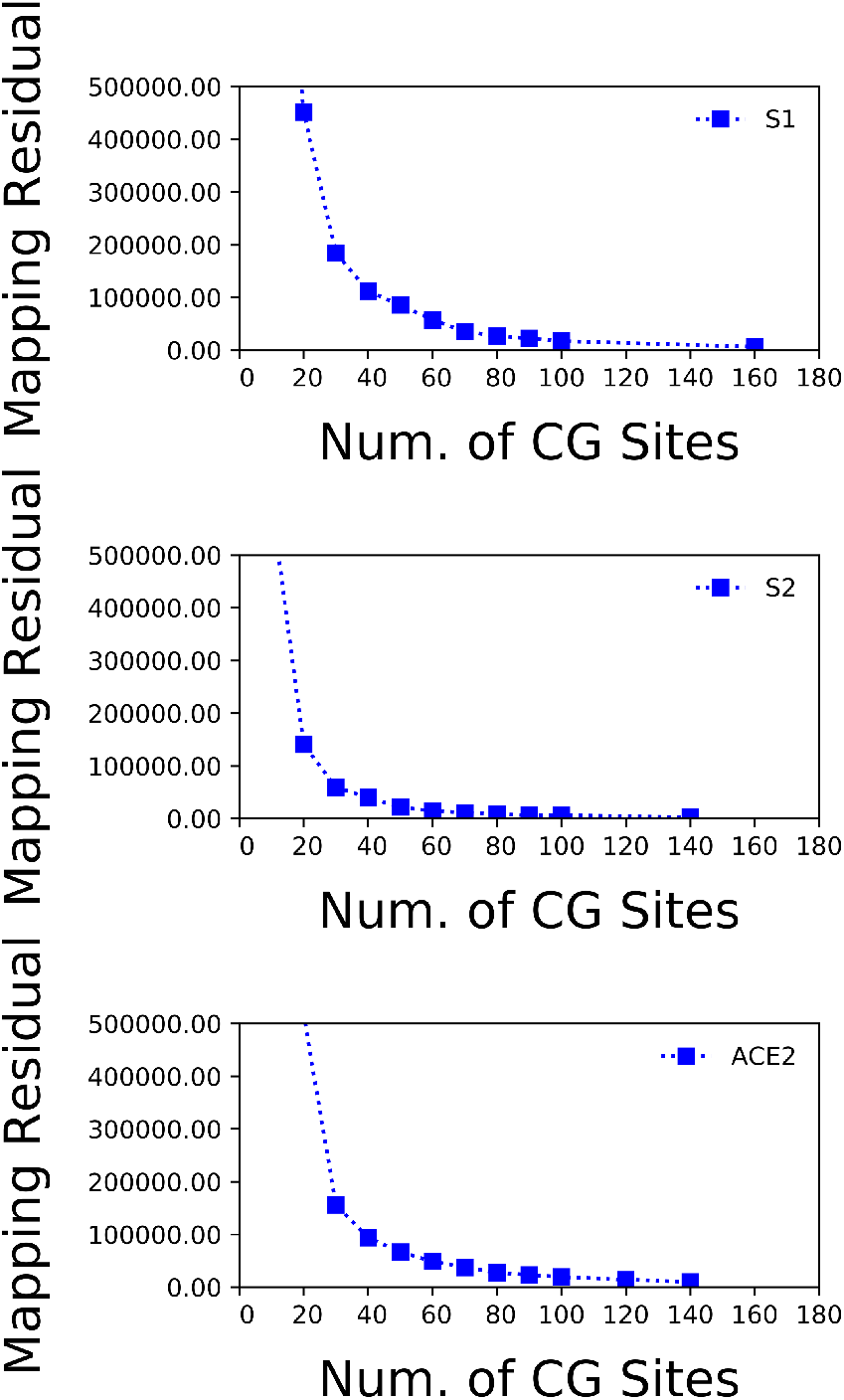
Comparison of EDCG residuals to CG model resolutions for each of the CG protein models: S1, S2, and ACE2.

The second phase determined the effective CG Hamiltonian for each protein. We systematically determined parameters for S1/S2 in the open state, S1/S2 in the closed state, and ACE2 in the bound state, as well as their inter-domain interactions. The Hamiltonian (*E*_*tot*_) was decomposed into four terms: intra-domain fluctuations (*E*_*intra*_), inter-domain exclusion (*E*_*excl*_), inter-domain attraction (*E*_*attr*_), and screened electrostatics (*E*_*yukawa*_):

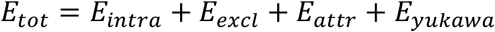

We used a hetero-elastic network model (hENM), in which harmonic bonds are assigned to all pairs of CG particles within a tunable distance cutoff *r*_*cut*_ (Lyman, Pfaendtner, & Voth, 2008), to represent *E*_*intra*_; bond energies are equal to *k*(*r* − *r*_0_)^2^ where *k* is the spring constant of a particular bond and *r*_0_ is the equilibrium bond length. The harmonic force constants *k* are iteratively adjusted until the fluctuations in the CG model converge to that of the atomistic data, i.e.,

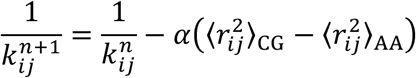

where 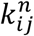 is the harmonic force constant for each *i*, *j* CG pair at iteration *n*, 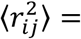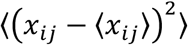 is the mean-squared fluctuation for each *i*, *j* CG pair, and *α* is a parameter that controls the magnitude of the adjustment for each iteration. We chose *r*_*cut*_ for each CG model using the “elbow” heuristic as shown in **Fig. 5**. Here, our residual was 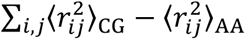. Following this protocol, we used distance cutoffs of 3, 4, and 3 nm for the S1, S2, and ACE2 CG models, respectively.

**Fig. 5.**
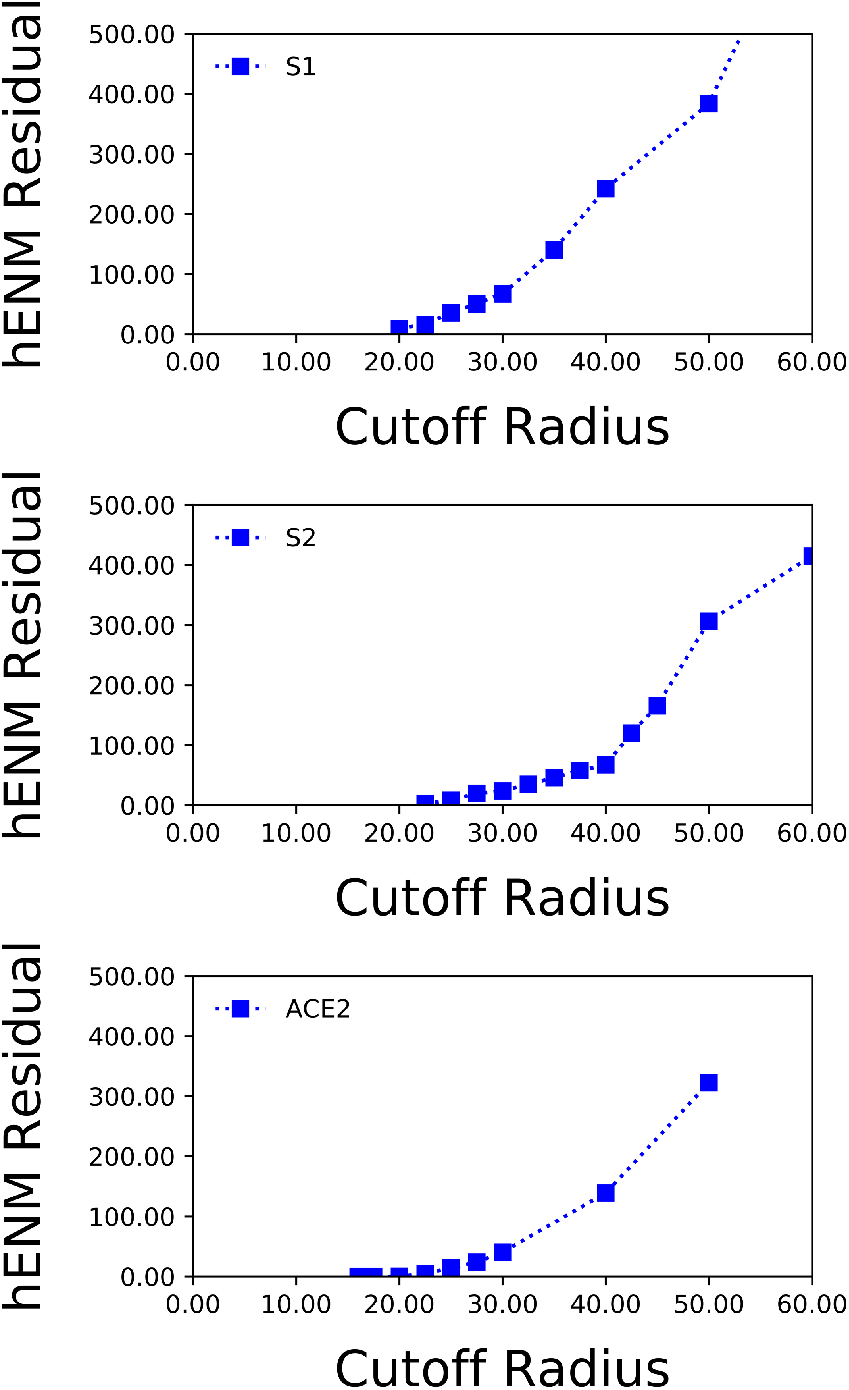
Comparison of hENM residuals to hENM cutoff radii for each of the CG protein models: S1, S2, and ACE2.

Inter-protein interactions were composed of excluded volume, attractive, and screened electrostatic terms. For excluded volume interactions (*E*_*excl*_), a soft cosine potential was used:

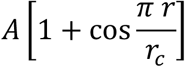

where *A* = 25 kcal/mol and *r*_*c*_ is the onset for excluded volume. For each CG pair, *r*_*c*_ was set to the minimum of two values: the minimum distance with non-zero frequency in the corresponding pairwise distance histogram from CG-mapped atomistic statistics or the default value of *r*_*c*_ = 3.0 nm. Screened electrostatics (*E*_*yukawa*_) were modeled using Yukawa potentials:

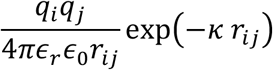

where *q*_*i*_ is the aggregate charge of CG site *i* κ = 1.274 nm^−1^ is the inverse Debye length for 0.15M NaCl, and ∊_*r*_ is the effective dielectric constant of the protein environment, approximated as 17.5 (Li, Li, Zhang, & Alexov, 2013).

Attractive non-bonded interactions between inter-protein contacts (*E*_*attr*_) were modeled as the sum of two Gaussian potentials:

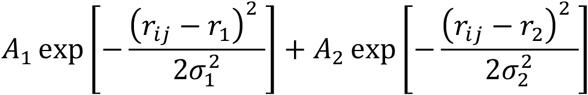

where *r*_1_ and σ_1_ are the mean and standard deviation determined by a fit to the pair correlation functions (from CG-mapped atomistic statistics) between CG sites *i* and *j* through least-squares regression. The constants *A*_1_ and *A*_2_ were optimized using relative-entropy minimization (REM) (Shell, 2008). We used the iterative Newton-Raphson method (Chaimovich & Shell, 2009) to update *A*_1_ and *A*_2_, which we refer to as parameter λ at iteration *n*:

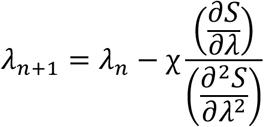

where χ is the “mixing ratio” or “learning rate” of the iterative optimization procedure. Each iteration required a LAMMPS simulation (Plimpton, 1995) of a single copy of the CG protein using the current CG force field, which was run for 21×10^6^ steps using a 100 fs timestep and a Langevin thermostat (10 ps damping time) at 300 K. Statistics were gathered every 800 steps over the final 20×10^6^ steps to calculate the parameter update. To aid convergence during training, a learning rate schedule for χ was implemented:

1. *Χ* = 0.5 during the first 25 iterations
2. *Χ* = 0.1 during the next 75 iterations
3. *Χ* = 0.01 during the next 75 iterations
4. *Χ* = 0.001 during the next 75 iterations
5. *Χ* = 0.1 during the next 50 iterations
6. *Χ* = 0.01 during the final 150 iterations

A total of 450 iterations were used to generate each CG model. After this point, changes to λ were effectively zero, and the optimization was considered complete.

The third phase determined an effective Hamiltonian for the spike trimer that was capable of continuously transitioning between the open and closed state. Here, we modified *E*_*intra*_, *E*_*excl*_, and *E*_*attr*_ as follows. For *E*_*intra*_, we kept bonds that appeared in both open and closed states and used the mean *k* and *r*_0_ constants. For *E*_*excl*_, we used the minimum *r*_*c*_ from both open and closed states. For *E*_*attr*_, we tested different linear combinations of the Gaussians parametrized from the open and closed state. We found that an equal mix recapitulated open/closed statistics the best, i.e., *E*_*attr*_ = 0.5*E*_*attr,open*_ + 0.5*E*_*attr,closed*_. No modification of *E*_*yukawa*_ was needed since this term is not state dependent.

The lipid model used the functional form from (Grime & Madsen, 2019), which we summarize here. We used a linear four-bead model with 1 head bead, 1 middle bead, and 2 tail beads. A piecewise potential was used to describe inter-lipid interactions:

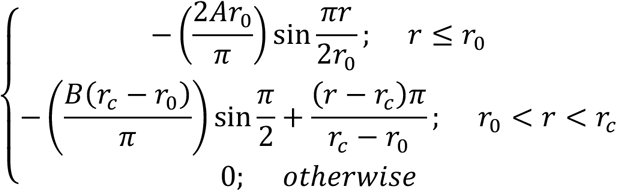

where *r*_0_ = 1.2 nm (or *r*_0_ = 0.9 nm for the head bead), *r*_*c*_ = 2.4 nm, *B* = 1.66 *k*_*B*_*T*, and *A* = 50 *k*_*B*_*T* (20 *k*_*B*_*T* for the head bead). Harmonic bonds and angles were used to describe intra-lipid interactions:

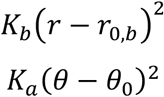

where *K*_b_ = 5 *k*_*B*_*T*/Å^2^, *r*_0,b_ = 0.6 nm, *K*_*a*_ = 5 *k*_*B*_*T*/rad^2^, and *θ*_0_ = *π* rad. For the purposes of modeling the interaction between the transmembrane domains of the spike trimer/ACE2 dimer and lipids, we effectively considered the transmembrane domain as having the same interaction as the lipids.

Model parameters and input files are available on Github: https://github.com/uchicago-voth/MSCG-models/tree/master/Spike_ACE2

### CG simulations

All simulations were prepared using PACKMOL and Moltemplate (Jewett, Zhuang, & Shea, 2013; Martínez, Andrade, Birgin, & Martínez, 2009). Briefly, CG spike trimers and ACE2 dimers were arranged in a grid using each density described in the main text. Two CG lipid membranes were randomly packed around the transmembrane domains of each grid of proteins in a 200×200 nm^2^ lateral (*xy*) domain; the *z* dimension of the box was fixed to 160 nm. The two membranes were then translated along the *z* direction until the minimum *z* distance between spike trimers and ACE2 dimers was 2 nm. All simulations were performed using LAMMPS (Plimpton, 1995). Conjugate gradient energy minimization was performed on each system until the change in force was less than 10^−6^. Then, proteins were held fixed while lipids were integrated using a Langevin thermostat at 310 K (0.5 ps damping constant) and a Berendsen barostat at 1 atm (0.5 ps damping constant, applied over the lateral *xy* dimension) over 2×10^6^ τ_CG_ (with τ_CG_ = 50 fs). During this process, the *xy* simulation domain compressed to around 170×170 nm^2^. We then integrated both proteins and lipids using a Langevin thermostat at 310 K (5 ps damping constant) over 170×10^6^ *τ*_CG_. Trajectory snapshots were saved every 25×10^3^ τ_CG_.

### Analysis

All analysis was performed using a combination of custom Tcl scripts using VMD (Humphrey, Dalke, & Schulten, 1996) and Python scripts using the MSMBuilder (Beauchamp et al., 2011) and MDTraj (McGibbon et al., 2015) libraries. The configurational state of each RBD was featurized using the DRID method (T. Zhou & Caflisch, 2012). Dimensional reduction (via tICA (Naritomi & Fuchigami, 2011)) of this feature set into 10 tICs was performed using a lag time of 40×10^3^ τ_CG_; the values of the first tIC eigenvector are shown in **Fig. 1 – figure supplement 1**. The data were then clustered into 3 sets using k-means clustering (Lloyd, 1982). We featurized and projected the configuration of each RBD extracted from our CG simulations onto the first tIC to classify its state. We also used the following distance metric to classify the bound state of the S1 domain to both its partner S2 domain and ACE2:

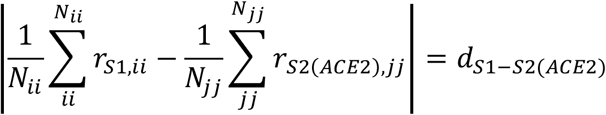

where *ii* and *jj* refer to CG site indices out of a set with *N*_*ii*_ and *N*_*jj*_ total sites, *r* is the position, *d* is the distance, and | | is the norm of the vector. In other words, we compute the distance between the centers of geometry of sets of CG sites. We chose CG site clusters that were observed to have close association distances with minimal fluctuation during CG model training. For S1/S2 binding, these included site indices 31-32 and 54-57 for S1 and 77, 87-89, 98-99, and 102 for S2. For S1/ACE2 binding, these included site indices 43-49 for S1 and 134-137, 158, and 162 for ACE2 (residue mapping for each site is shown in Supplementary File 1). Using this metric, we respectively classified S1/S2 and S1/ACE2 as bound states when *d*_*S1-S2*_ and *d*_*S1-ACE2*_ were less than 4 nm, respectively.

## Acknowledgements

This work was supported in part by the National Science Foundation through NSF RAPID grant CHE-2029092 (A.J.P and G.A.V). This study was also supported by funding from the European Research Council (ERC) under the European Union's Horizon 2020 research and innovation programme (ERC-CoG-648432 MEMBRANEFUSION to J.A.G.B.) and the Medical Research Council as part of UK Research and Innovation (MC_UP_1201/16 to J.A.G.B.). Alexander J. Pak gratefully acknowledges support from the National Institute of Allergy and Infectious Diseases of the National Institutes of Health under grant F32 AI150477. Alvin Yu gratefully acknowledges support from the National Institute of Allergy and Infectious Diseases of the National Institutes of Health under grant F32 AI150208. This work used resources, services, and support provided via the COVID-19 HPC Consortium (https://covid19-hpc-consortium.org/), which is a unique private-public effort to bring together government, industry, and academic leaders who are volunteering free compute time and resources in support of COVID-19 research. The authors acknowledge the use of the Witherspoon cluster at IBM Research that has contributed to this work. This research also used resources of the Oak Ridge Leadership Computing Facility at the Oak Ridge National Laboratory, which is supported by the Office of Science of the U.S. Department of Energy under Contract No. DE-AC05-00OR22725. The authors also acknowledge the Texas Advanced Computing Center (TACC) at The University of Texas at Austin for providing HPC and visualization resources that have contributed to the research results reported within this paper.

## Competing Interests

The authors declare no competing financial and non-financial interests.

## Supplementary Files

**Supplementary File 1** – Tables that summarize the residue mapping to each coarse-grained site type in S1, S2, and ACE2.

## Figure Supplements

**Fig. 1 - figure supplement 1.**
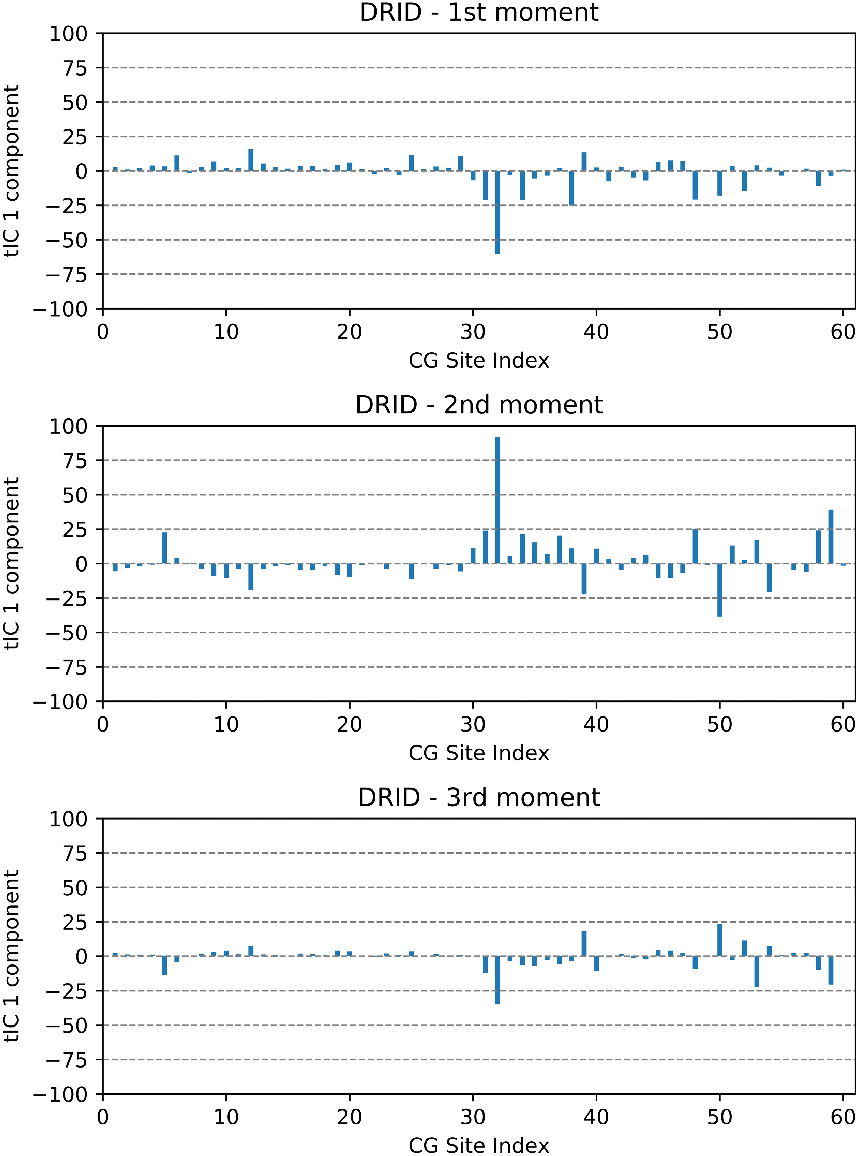
Eigenvector components of the 1^st^ tIC with respect to the 1^st^ through 3^rd^ moments of DRID using the listed CG bead index within S1 as the reference point.

**Fig. 1 - figure supplement 2.**
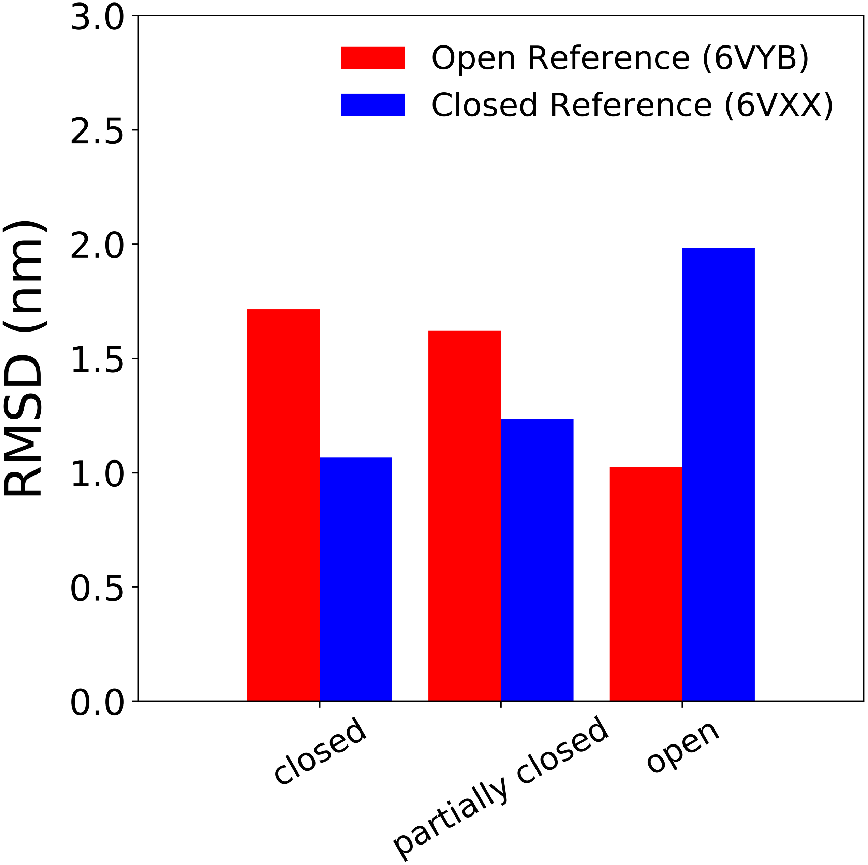
Root-mean-squared deviation (RMSD) of coarse-grained S1 configurations from the closed, partially closed, and open k-means clusters using the closed and open configurations from PDB 6VXX and 6VYB, respectively, as reference (Walls et al., 2020).

**Fig. 2 - figure supplement 1.**
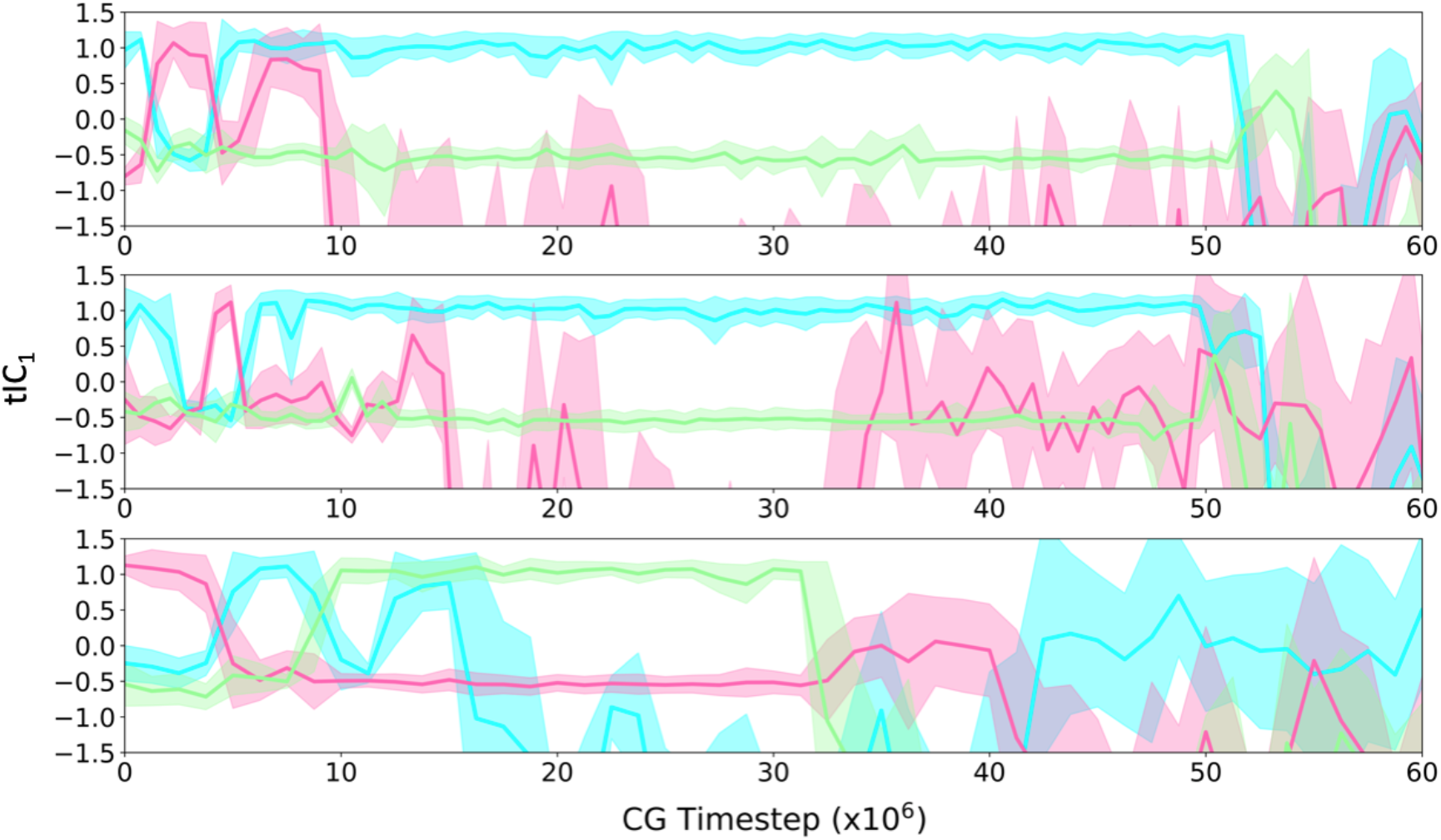
Additional time-series profiles of tIC_1_ during ACE2-mediated S1 dissociation from spike trimers. Each of the three panels is an independent spike trimer. The colors are the same as **Fig. 2**. Note that the spike trimer protomers are labeled cyan to pink to green to cyan in counter-clockwise order when viewing from the top-down. These time series profiles indicate that the counter-clockwise protomer from the initially ACE2-bound protomer is bound next then dissociated.

**Video 1.**
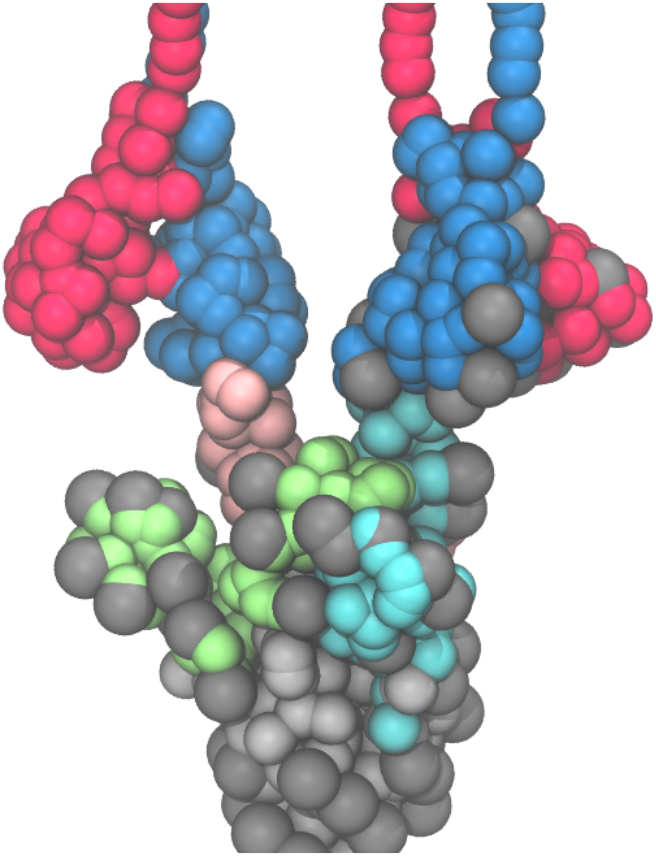
Dissociation of S1 induced by multivalent ACE2 binding during coarse-grained molecular dynamics simulations. The two protomers within ACE2 dimers are depicted as red and blue beads, respectively, while the three S1 protomers within the spike trimer are depicted as cyan, pink, and green beads. All glycans and S2 protomers are represented by grey and silver beads, respectively. Lipids are not shown for clarity.

## Notes

### Competing Interest Statement

The authors have declared no competing interest.

### Summary of Updates

New Preprint Version

## References

Abraham, M. J., Murtola, T., Schulz, R., Páll, S., Smith, J. C., Hess, B., & Lindahl, E. (2015). GROMACS: High performance molecular simulations through multi-level parallelism from laptops to supercomputers. SoftwareX, 1-2, 19–25. doi:https://doi.org/10.1016/j.softx.2015.06.001

Anand, S. P., Chen, Y., Prévost, J., Gasser, R., Beaudoin-Bussières, G., Abrams, C. F., … Finzi, A. (2020). Interaction of Human ACE2 to Membrane-Bound SARS-CoV-1 and SARS-CoV-2 S Glycoproteins. Viruses, 12(10). doi:10.3390/v12101104

Barros, E. P., Casalino, L., Gaieb, Z., Dommer, A. C., Wang, Y., Fallon, L., … Amaro, R. E. The Flexibility of ACE2 in the Context of SARS-CoV-2 Infection. Biophysical Journal, 120, 1072–1084. Retrieved from https://doi.org/10.1016/j.bpj.2020.10.036

Beauchamp, K. A., Bowman, G. R., Lane, T. J., Maibaum, L., Haque, I. S., & Pande, V. S. (2011). MSMBuilder2: Modeling Conformational Dynamics on the Picosecond to Millisecond Scale. Journal of Chemical Theory and Computation, 7(10), 3412–3419. doi:10.1021/ct200463m

Benton, D. J., Wrobel, A. G., Roustan, C., Borg, A., Xu, P., Martin, S. R., … Gamblin, S. J. (2021). The effect of the D614G substitution on the structure of the spike glycoprotein of SARS-CoV-2. Proceedings of the National Academy of Sciences, 118(9), e2022586118. doi:10.1073/pnas.2022586118

Benton, D. J., Wrobel, A. G., Xu, P., Roustan, C., Martin, S. R., Rosenthal, P. B., … Gamblin, S. J. (2020). Receptor binding and priming of the spike protein of SARS-CoV-2 for membrane fusion. Nature, 588(7837), 327–330. doi:10.1038/s41586-020-2772-0

Bussi, G., Zykova-Timan, T., & Parrinello, M. (2009). Isothermal-isobaric molecular dynamics using stochastic velocity rescaling. The Journal of Chemical Physics, 130(7), 074101. doi:10.1063/1.3073889

Cai, Y., Zhang, J., Xiao, T., Peng, H., Sterling, S. M., Walsh, R. M., … Chen, B. (2020). Distinct conformational states of SARS-CoV-2 spike protein. Science, 369(6511), 1586. doi:10.1126/science.abd4251

Casalino, L., Dommer, A. C., Gaieb, Z., Barros, E. P., Sztain, T., Ahn, S.-H., … Amaro, R. E. (2021). AI-driven multiscale simulations illuminate mechanisms of SARS-CoV-2 spike dynamics. The International Journal of High Performance Computing Applications, 10943420211006452. doi:10.1177/10943420211006452

Casalino, L., Gaieb, Z., Goldsmith, J. A., Hjorth, C. K., Dommer, A. C., Harbison, A. M., … Amaro, R. E. (2020). Beyond Shielding: The Roles of Glycans in the SARS-CoV-2 Spike Protein. ACS Central Science, 6(10), 1722–1734. doi:10.1021/acscentsci.0c01056

Chaimovich, A., & Shell, M. S. (2009). Anomalous waterlike behavior in spherically-symmetric water models optimized with the relative entropy. Physical Chemistry Chemical Physics, 11(12), 1901–1915. doi:10.1039/B818512C

Grime, J. M. A., & Madsen, J. J. (2019). Efficient Simulation of Tunable Lipid Assemblies Across Scales and Resolutions. arXiv:1910.05362v1.

Gur, M., Taka, E., Yilmaz, S. Z., Kilinc, C., Aktas, U., & Golcuk, M. (2020). Conformational transition of SARS-CoV-2 spike glycoprotein between its closed and open states. The Journal of Chemical Physics, 153(7), 075101. doi:10.1063/5.0011141

Hess, B., Bekker, H., Berendsen, H. J. C., & Fraaije, J. G. E. M. (1997). LINCS: A linear constraint solver for molecular simulations. Journal of Computational Chemistry, 18(12), 1463–1472. doi:10.1002/(SICI)1096-987X(199709)18:12<1463::AID-JCC4>3.0.CO;2-H</1463::AID-JCC4>

Huang, J., Rauscher, S., Nawrocki, G., Ran, T., Feig, M., de Groot, B. L., … MacKerell, A. D. Jr (2016). CHARMM36m: an improved force field for folded and intrinsically disordered proteins. Nature Methods, 14, 71. doi:10.1038/nmeth.4067 https://www.nature.com/articles/nmeth.4067#supplementary-information

Humphrey, W., Dalke, A., & Schulten, K. (1996). VMD: Visual molecular dynamics. Journal of Molecular Graphics, 14(1), 33–38. doi:https://doi.org/10.1016/0263-7855(96)00018-5

Jewett, A. I., Zhuang, Z., & Shea, J.-E. (2013). Moltemplate a Coarse-Grained Model Assembly Tool. Biophysical Journal, 104(2), 169a. doi:10.1016/j.bpj.2012.11.953

Jo, S., Kim, T., Iyer, V. G., & Im, W. (2008). CHARMM-GUI: A web-based graphical user interface for CHARMM. Journal of Computational Chemistry, 29(11), 1859–1865. doi:https://doi.org/10.1002/jcc.20945

Ke, Z., Oton, J., Qu, K., Cortese, M., Zila, V., McKeane, L., … Briggs, J. A. G. (2020). Structures and distributions of SARS-CoV-2 spike proteins on intact virions. Nature, 588(7838), 498–502. doi:10.1038/s41586-020-2665-2

Li, L., Li, C., Zhang, Z., & Alexov, E. (2013). On the Dielectric “Constant” of Proteins: Smooth Dielectric Function for Macromolecular Modeling and Its Implementation in DelPhi. Journal of Chemical Theory and Computation, 9(4), 2126–2136. doi:10.1021/ct400065j

Lloyd, S. (1982). Least squares quantization in PCM. IEEE Transactions on Information Theory, 28(2), 129–137. doi:10.1109/TIT.1982.1056489

Lu, M., Uchil, P. D., Li, W., Zheng, D., Terry, D. S., Gorman, J., … Mothes, W. (2020). Real-Time Conformational Dynamics of SARS-CoV-2 Spikes on Virus Particles. Cell Host & Microbe, 28(6), 880–891.e888. doi:https://doi.org/10.1016/j.chom.2020.11.001

Lui, I., Zhou, X. X., Lim, S. A., Elledge, S. K., Solomon, P., Rettko, N. J., … Wells, J. A. (2020). Trimeric SARS-CoV-2 Spike interacts with dimeric ACE2 with limited intra-Spike avidity. bioRxiv. doi:10.1101/2020.05.21.109157

Lyman, E., Pfaendtner, J., & Voth, G. A. (2008). Systematic Multiscale Parameterization of Heterogeneous Elastic Network Models of Proteins. Biophysical Journal, 95(9), 4183–4192. doi:10.1529/biophysj.108.139733

Mansbach, R. A., Chakraborty, S., Nguyen, K., Montefiori, D. C., Korber, B., & Gnanakaran, S. (2021). The SARS-CoV-2 Spike variant D614G favors an open conformational state. Science Advances, 7(16), eabf3671. doi:10.1126/sciadv.abf3671

Maragakis, P., & Karplus, M. (2005). Large Amplitude Conformational Change in Proteins Explored with a Plastic Network Model: Adenylate Kinase. Journal of Molecular Biology, 352(4), 807–822. doi:https://doi.org/10.1016/j.jmb.2005.07.031

Martínez, L., Andrade, R., Birgin, E. G., & Martínez, J. M. (2009). PACKMOL: A package for building initial configurations for molecular dynamics simulations. Journal of Computational Chemistry, 30(13), 2157–2164. doi:https://doi.org/10.1002/jcc.21224

Martyna, G. J., Klein, M. L., & Tuckerman, M. (1992). Nosé–Hoover chains: The canonical ensemble via continuous dynamics. The Journal of Chemical Physics, 97(4), 2635–2643. doi:10.1063/1.463940

McGibbon Robert, T., Beauchamp Kyle, A., Harrigan Matthew, P., Klein, C., Swails Jason, M., Hernández Carlos, X., … Pande Vijay, S. (2015). MDTraj: A Modern Open Library for the Analysis of Molecular Dynamics Trajectories. Biophysical Journal, 109(8), 1528–1532. doi:https://doi.org/10.1016/j.bpj.2015.08.015

Naritomi, Y., & Fuchigami, S. (2011). Slow dynamics in protein fluctuations revealed by time-structure based independent component analysis: The case of domain motions. The Journal of Chemical Physics, 134(6), 065101. doi:10.1063/1.3554380

Ozono, S., Zhang, Y., Ode, H., Sano, K., Tan, T. S., Imai, K., … Tokunaga, K. (2021). SARS-CoV-2 D614G spike mutation increases entry efficiency with enhanced ACE2-binding affinity. Nature Communications, 12(1), 848. doi:10.1038/s41467-021-21118-2

Parrinello, M., & Rahman, A. (1980). Crystal Structure and Pair Potentials: A Molecular-Dynamics Study. Physical Review Letters, 45(14), 1196–1199. doi:10.1103/PhysRevLett.45.1196

Plimpton, S. (1995). Fast Parallel Algorithms for Short-Range Molecular Dynamics. Journal of Computational Physics, 117(1), 1–19. doi:https://doi.org/10.1006/jcph.1995.1039

Raghuvamsi, P. V., Tulsian, N. K., Samsudin, F., Qian, X., Purushotorman, K., Yue, G., … Anand, G. S. (2021). SARS-CoV-2 S protein:ACE2 interaction reveals novel allosteric targets. eLife, 10, e63646. doi:10.7554/eLife.63646

Shang, J., Wan, Y., Luo, C., Ye, G., Geng, Q., Auerbach, A., & Li, F. (2020). Cell entry mechanisms of SARS-CoV-2. Proceedings of the National Academy of Sciences, 117(21), 11727. doi:10.1073/pnas.2003138117

Sharp, M. E., Vázquez, F. X., Wagner, J. W., Dannenhoffer-Lafage, T., & Voth, G. A. (2019). Multiconfigurational Coarse-Grained Molecular Dynamics. Journal of Chemical Theory and Computation, 15(5), 3306–3315. doi:10.1021/acs.jctc.8b01133

Shell, M. S. (2008). The relative entropy is fundamental to multiscale and inverse thermodynamic problems. The Journal of Chemical Physics, 129(14), 144108. doi:10.1063/1.2992060

Supasa, P., Zhou, D., Dejnirattisai, W., Liu, C., Mentzer, A. J., Ginn, H. M., … Screaton, G. R. (2021). Reduced neutralization of SARS-CoV-2 B.1.1.7 variant by convalescent and vaccine sera. Cell, 184(8), 2201–2211.e2207. doi:https://doi.org/10.1016/j.cell.2021.02.033

Turoňová, B., Sikora, M., Schürmann, C., Hagen, W. J. H., Welsch, S., Blanc, F. E. C., … Beck, M. (2020). In situ structural analysis of SARS-CoV-2 spike reveals flexibility mediated by three hinges. Science, 370(6513), 203. doi:10.1126/science.abd5223

Walls, A. C., Park, Y.-J., Tortorici, M. A., Wall, A., McGuire, A. T., & Veesler, D. (2020). Structure, Function, and Antigenicity of the SARS-CoV-2 Spike Glycoprotein. Cell, 183, 281–292. Retrieved from http://www.ncbi.nlm.nih.gov/pubmed/32155444

Walls, A. C., Tortorici, M. A., Snijder, J., Xiong, X., Bosch, B.-J., Rey, F. A., & Veesler, D. (2017). Tectonic conformational changes of a coronavirus spike glycoprotein promote membrane fusion. Proceedings of the National Academy of Sciences, 114(42), 11157. doi:10.1073/pnas.1708727114

Watanabe, Y., Allen, J. D., Wrapp, D., McLellan, J. S., & Crispin, M. (2020). Site-specific glycan analysis of the SARS-CoV-2 spike. Science, 369, 330–333. Retrieved from http://science.sciencemag.org/content/early/2020/05/01/science.abb9983.abstract

Wrapp, D., Wang, N., Corbett, K. S., Goldsmith, J. A., Hsieh, C.-L., Abiona, O., … McLellan, J. S. (2020). Cryo-EM structure of the 2019-nCoV spike in the prefusion conformation. Science, 367(6483), 1260. doi:10.1126/science.abb2507

Xiao, T., Lu, J., Zhang, J., Johnson, R. I., McKay, L. G. A., Storm, N., … Chen, B. (2021). A trimeric human angiotensin-converting enzyme 2 as an anti-SARS-CoV-2 agent. Nature Structural & Molecular Biology, 28(2), 202–209. doi:10.1038/s41594-020-00549-3

Xu, C., Wang, Y., Liu, C., Zhang, C., Han, W., Hong, X., … Cong, Y. (2021). Conformational dynamics of SARS-CoV-2 trimeric spike glycoprotein in complex with receptor ACE2 revealed by cryo-EM. Science Advances, 7(1), eabe5575. doi:10.1126/sciadv.abe5575

Yan, R., Zhang, Y., Li, Y., Xia, L., Guo, Y., & Zhou, Q. (2020). Structural basis for the recognition of SARS-CoV-2 by full-length human ACE2. Science, 367(6485), 1444. doi:10.1126/science.abb2762

Yang, J., Petitjean, S. J. L., Koehler, M., Zhang, Q., Dumitru, A. C., Chen, W., … Alsteens, D. (2020). Molecular interaction and inhibition of SARS-CoV-2 binding to the ACE2 receptor. Nature Communications, 11(1), 4541. doi:10.1038/s41467-020-18319-6

Yu, A., Pak, A. J., He, P., Monje-Galvan, V., Casalino, L., Gaieb, Z., … Voth, G. A. (2020). A Multiscale Coarse-Grained Model of the SARS-CoV-2 Virion. Biophysical Journal, 120, 1097–1104. Retrieved from http://www.sciencedirect.com/science/article/pii/S0006349520331684

Zhang, L., Jackson, C. B., Mou, H., Ojha, A., Peng, H., Quinlan, B. D., … Choe, H. (2020). SARS-CoV-2 spike-protein D614G mutation increases virion spike density and infectivity. Nature Communications, 11(1), 6013. doi:10.1038/s41467-020-19808-4

Zhang, Z., Lu, L., Noid, W. G., Krishna, V., Pfaendtner, J., & Voth, G. A. (2008). A Systematic Methodology for Defining Coarse-Grained Sites in Large Biomolecules. Biophysical Journal, 95(11), 5073–5083. doi:10.1529/biophysj.108.139626

Zhou, D., Dejnirattisai, W., Supasa, P., Liu, C., Mentzer, A. J., Ginn, H. M., … Screaton, G. R. (2021). Evidence of escape of SARS-CoV-2 variant B.1.351 from natural and vaccine-induced sera. Cell, 184(9), 2348–2361.e2346. doi:10.1016/j.cell.2021.02.037

Zhou, T., & Caflisch, A. (2012). Distribution of Reciprocal of Interatomic Distances: A Fast Structural Metric. Journal of Chemical Theory and Computation, 8(8), 2930–2937. doi:10.1021/ct3003145

