## Supplemental File for "Cooperative multivalent receptor binding promotes exposure of the SARS-CoV-2 fusion machinery core"

#### Supplementary File 1 for

### Cooperative multivalent receptor binding promotes exposure of the SARS-CoV-2 fusion machinery core

Alexander J. Pak[1,⊥], Alvin Yu[1], Zunlong Ke[2], John A. G. Briggs[2], and Gregory A. Voth[1,#]

[1] Department of Chemistry, Chicago Center for Theoretical Chemistry, Institute for Biophysical Dynamics, and James Franck Institute, The University of Chicago, Chicago, IL, USA

[2] Structural Studies Division, Medical Research Council Laboratory of Molecular Biology, Cambridge, UK

[⊥] Present address: Department of Chemical and Biological Engineering, Colorado School of Mines, Golden, CO 80401

Supplementary File 1 includes **Tables S1-S3.**

**Table S1.** Summary of residue mapping to each coarse-grained site type in S1 (glycans (CG types 61-73) excluded).

**S1 Protein**

| CG Type | Residue (Start) | Residue (End) | CG Index | Residue (Start) | Residue (End) |
|---|---|---|---|---|---|
| 1 | 16 | 19 | 31 | 289 | 308 |
| 2 | 20 | 23 | 32 | 309 | 319 |
| 3 | 24 | 28 | 33 | 320 | 328 |
| 4 | 29 | 37 | 34 | 329 | 336 |
| 5 | 38 | 53 | 35 | 337 | 346 |
| 6 | 54 | 64 | 36 | 347 | 355 |
| 7 | 65 | 70 | 37 | 356 | 364 |
| 8 | 71 | 75 | 38 | 365 | 375 |
| 9 | 76 | 80 | 39 | 376 | 387 |
| 10 | 81 | 92 | 40 | 388 | 401 |
| 11 | 93 | 103 | 41 | 402 | 418 |
| 12 | 104 | 118 | 42 | 419 | 433 |
| 13 | 119 | 130 | 43 | 434 | 449 |
| 14 | 131 | 140 | 44 | 450 | 458 |
| 15 | 141 | 146 | 45 | 459 | 467 |
| 16 | 147 | 151 | 46 | 468 | 474 |

| | | | | | |
|---|---|---|---|---|---|
| 17 | 152 | 156 | 47 | 475 | 482 |
| 18 | 157 | 162 | 48 | 483 | 492 |
| 19 | 163 | 173 | 49 | 493 | 510 |
| 20 | 174 | 182 | 50 | 511 | 524 |
| 21 | 183 | 191 | 51 | 525 | 547 |
| 22 | 192 | 206 | 52 | 548 | 574 |
| 23 | 207 | 222 | 53 | 575 | 593 |
| 24 | 223 | 233 | 54 | 594 | 617 |
| 25 | 234 | 244 | 55 | 618 | 626 |
| 26 | 245 | 250 | 56 | 627 | 633 |
| 27 | 251 | 256 | 57 | 634 | 640 |
| 28 | 257 | 261 | 58 | 641 | 658 |
| 29 | 262 | 272 | 59 | 659 | 677 |
| 30 | 273 | 288 | 60 | 678 | 685 |

**Table S2.** Summary of residue mapping to each coarse-grained site type in S2 (glycans (CG types 124-132) excluded).

**S2 Protein**

| CG Type | Residue (Start) | Residue (End) | CG Index | Residue (Start) | Residue (End) |
|---|---|---|---|---|---|
| 74 | 686 | 689 | 99 | 958 | 970 |
| 75 | 690 | 697 | 100 | 971 | 984 |
| 76 | 698 | 705 | 101 | 985 | 999 |
| 77 | 706 | 718 | 102 | 1000 | 1014 |
| 78 | 719 | 726 | 103 | 1015 | 1028 |
| 79 | 727 | 736 | 104 | 1029 | 1051 |
| 80 | 737 | 754 | 105 | 1052 | 1065 |
| 81 | 755 | 768 | 106 | 1066 | 1077 |
| 82 | 767 | 781 | 107 | 1078 | 1093 |
| 83 | 782 | 791 | 108 | 1094 | 1115 |
| 84 | 792 | 806 | 109 | 1116 | 1137 |
| 85 | 807 | 813 | 110 | 1138 | 1149 |
| 86 | 814 | 828 | 111 | 1150 | 1160 |
| 87 | 829 | 835 | 112 | 1161 | 1167 |
| 88 | 836 | 840 | 113 | 1168 | 1175 |
| 89 | 841 | 845 | 114 | 1176 | 1186 |
| 90 | 846 | 851 | 115 | 1187 | 1196 |
| 91 | 852 | 861 | 116 | 1197 | 1206 |
| 92 | 862 | 875 | 117 | 1207 | 1214 |
| 93 | 876 | 893 | 118 | 1215 | 1228 |
| 94 | 894 | 910 | 119 | 1229 | 1243 |
| 95 | 911 | 923 | 120 | 1244 | 1251 |
| 96 | 924 | 935 | 121 | 1252 | 1258 |
| 97 | 936 | 944 | 122 | 1259 | 1267 |
| 98 | 945 | 957 | 123 | 1268 | 1273 |

**Table S3.** Summary of residue mapping to each coarse-grained site type in ACE2 (glycans (CG types 203-209) excluded).

**ACE2 Protein**

| CG Type | Residue (Start) | Residue (End) | CG Index | Residue (Start) | Residue (End) |
|---|---|---|---|---|---|
| 133 | 21 | 31 | 168 | 425 | 430 |
| 134 | 32 | 42 | 169 | 431 | 441 |
| 135 | 43 | 53 | 170 | 442 | 454 |
| 136 | 54 | 63 | 171 | 455 | 467 |
| 137 | 64 | 73 | 172 | 468 | 478 |
| 138 | 74 | 83 | 173 | 479 | 490 |
| 139 | 84 | 94 | 174 | 491 | 499 |
| 140 | 94 | 103 | 175 | 500 | 510 |
| 141 | 104 | 108 | 176 | 511 | 523 |
| 142 | 109 | 118 | 177 | 524 | 535 |
| 143 | 119 | 132 | 178 | 536 | 543 |
| 144 | 133 | 140 | 179 | 544 | 554 |
| 145 | 141 | 154 | 180 | 555 | 565 |
| 146 | 155 | 166 | 181 | 566 | 577 |
| 147 | 167 | 182 | 182 | 578 | 588 |
| 148 | 183 | 193 | 183 | 589 | 598 |
| 149 | 194 | 204 | 184 | 599 | 605 |
| 150 | 205 | 214 | 185 | 606 | 611 |
| 151 | 215 | 226 | 186 | 612 | 619 |
| 152 | 227 | 244 | 187 | 620 | 625 |
| 153 | 245 | 264 | 188 | 626 | 629 |
| 154 | 265 | 281 | 189 | 630 | 633 |
| 155 | 282 | 292 | 190 | 634 | 643 |
| 156 | 293 | 304 | 191 | 644 | 653 |
| 157 | 305 | 320 | 192 | 654 | 663 |
| 158 | 321 | 332 | 193 | 664 | 672 |
| 159 | 333 | 336 | 194 | 673 | 684 |
| 160 | 337 | 340 | 195 | 685 | 705 |
| 161 | 341 | 345 | 196 | 706 | 724 |
| 162 | 346 | 357 | 197 | 725 | 732 |
| 163 | 358 | 363 | 198 | 733 | 739 |
| 164 | 364 | 378 | 199 | 740 | 747 |
| 165 | 379 | 394 | 200 | 748 | 754 |
| 166 | 395 | 409 | 201 | 755 | 764 |
| 167 | 410 | 424 | 202 | 765 | 768 |
